# Evolutionary sparse learning with paired species contrast reveals the shared genetic basis of convergent traits

**DOI:** 10.1101/2025.01.08.631987

**Authors:** John B. Allard, Sudip Sharma, Ravi Patel, Maxwell Sanderford, Koichiro Tamura, Slobodan Vucetic, Glenn S. Gerhard, Sudhir Kumar

## Abstract

Cases abound in which nearly identical traits have appeared in distant species facing similar environments. These unmistakable examples of adaptive evolution offer opportunities to gain insight into their genetic origins and mechanisms through comparative analyses. Here, we present a novel comparative genomics approach to build genetic models that underlie the independent origins of convergent traits using evolutionary sparse learning. We test the hypothesis that common genes and sites are involved in the convergent evolution of two key traits: C4 photosynthesis in grasses and echolocation in mammals. Genetic models were highly predictive of independent cases of convergent evolution of C4 photosynthesis. These results support the involvement of sequence substitutions in many common genetic loci in the evolution of convergent traits studied. Genes contributing to genetic models for echolocation were highly enriched for functional categories related to hearing, sound perception, and deafness (*P* < 10^−6^); a pattern that has eluded previous efforts applying standard molecular evolutionary approaches. We conclude that phylogeny-informed machine learning naturally excludes apparent molecular convergences due to shared species history, enhances the signal-to-noise ratio for detecting molecular convergence, and empowers the discovery of common genetic bases of trait convergences.

---

Organisms continuously adapt to their natural environment. Under similar environmental conditions, the same adaptations may evolve independently in clades across the tree of life. For example, the convergent evolution of the ability to echolocate in some bats and toothed whales is an example of adaptation brought on by major transitions to new environments requiring similar physiological innovations. Evolutionary biologists have long sought the common genetic basis of these convergent adaptations under the hypothesis that the same pathways, genes, and/or base substitutions are involved in these adaptations. However, “*the extent to which convergent traits evolve by similar genetic and molecular pathways is not clear*”^1^. Despite many molecular evolutionary investigations, the strongest evidence for molecular convergence thus far appears to be a marginally significant (FDR-corrected *P* = 0.0486) enrichment of sound perception genes in which convergent and parallel amino acid substitutions were observed^2–4^. Although these results hint at the possible presence of some shared genetic basis in the evolution of echolocation in independent clades, some studies could not detect such an enrichment^3^, casting doubt on the robustness of the results, the general applicability of the methodology, or even the presence of a common genetic basis.

The lack of consistent and statistically significant results may be due to insufficient commonality in the genetic bases of these traits, i.e., different genes and different sites may perform similar functions in independent clades. Alternatively, the lack of sufficient statistical power or inability to fully exclude non-adaptive convergence may be hampering efforts to detect genes and sites associated with the evolution of convergent traits^5–7^. Furthermore, current state-of-the-art approaches primarily reveal retrospective patterns, but they do not explicitly model quantitative genetic changes in convergent trait evolution to make statistical predictions of the presence or absence of the convergent trait.

We have addressed these challenges by building predictive genetic models of convergent trait evolution using evolutionary sparse learning (ESL). ESL is supervised machine learning in which genomic components (e.g., genes and sites) are model parameters, and substitutions in multiple sequence alignments are observations^8^. We developed a paired species contrast (PSC) design to select the training data for machine learning to automatically mask neutral (background) sequence convergence that can lead to spurious inferences and reduce the power to detect the genetic basis of convergence^5,6,9^. Importantly, ESL-PSC simultaneously considers all genetic loci and their respective substitutions during computational analysis, eliminating biases due to arbitrary evolutionary conservation thresholds and convergent substitution cut-offs necessary in some other approaches^2,3,7,10,11^.

ESL-PSC produces a quantitative genetic model to predict the presence/absence of a convergent trait in any species based on its genome sequence. This is needed to test the biological hypothesis of commonality of genetic basis in the independent evolution of the same trait. Lists of loci comprising the genetic model can be subjected to additional analysis to test if there is an enrichment of functional categories relevant to the trait analyzed^12,13^. This approach is commonly used to establish the biological relevance of candidate loci derived from large-scale scans for molecular convergence in the absence of alternatives^2–4,9,14–16^. We applied ESL-PSC to build genetic models of convergent evolution of C4 photosynthesis in grasses and of echolocation in mammals because they have been extensively investigated previously^4,17–22^.

## ESL-PSC for building genetic models of convergent traits

We introduce ESL-PSC with an analysis of protein sequence alignments of chloroplast proteins, which are well-suited for demonstrating the predictive ability of the method in a range of grass species that have acquired C4 photosynthesis independently. One may alternatively use ESL-PSC for nucleotide sequence alignments with the option to group sites into exons, introns, or other types of domains and functional annotations, as described in the *Material and Methods* section.

ESL uses logistic regression to infer a genetic model that can predict trait-positive and trait-negative species, which we numerically encode as +1 and −1, respectively^8,23^. In this analysis, the Least Absolute Shrinkage and Selection Operator (LASSO) compares alternative genetic models by imposing penalties for including additional amino acid positions and genes into the model while seeking high prediction accuracy. ESL-PSC produces models that incorporate only those proteins whose member sites make a significant contribution to the ability of the genetic model to classify species according to their traits rather than their ancestry.

To train the ESL model, we use a paired species contrast (PSC) approach in which a balanced training dataset of equal numbers of trait-positive and trait-negative species (those with and without the trait of interest, respectively) is first selected such that for every trait-positive species, we include one closely-related trait-negative species. In PSC, species pairs are required to be from evolutionarily independent clades to avoid introducing evolutionary correlations among pairs due to shared evolutionary history, which is known to cause spurious associations^5,6,9^. As an example, we could select trait-positive species A_1_ and D_1_ and trait-negative species B_1_ and C, respectively, to satisfy the above conditions (**Fig. 1A**).

**Figure 1.**
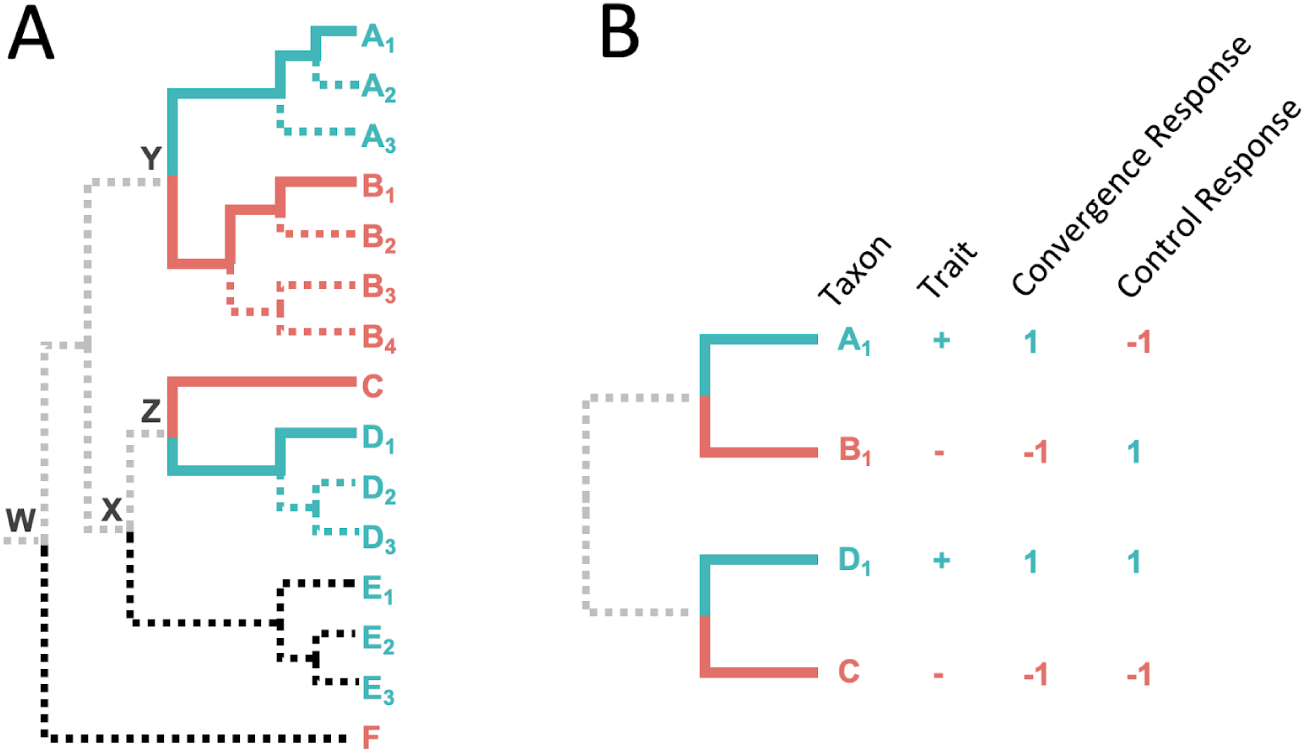
The paired species contrast (PSC) design. **A**: An example phylogeny with one set of selected species (solid blue and red lines). Extraneous lineages (black dotted lines) and shared evolutionary history (gray dotted lines). **B**: A schematic depiction of the four species selected for ESL-PSC analysis. In the ESL experiment, the response variable refers to the binary phenotype, where +1 represents the convergent trait, and −1 represents the ancestral trait.

PSC selection of training data ensures that the most recent common ancestor (MRCA) of each trait-positive and trait-negative species pair selected will be more recent than the MRCA of either member of the pair with any of the other species in the analysis. In the above example, the MRCA of A_1_ and B_1_ (Y) is more recent than that of A_1_ and F (W). Also, ESL-PSC automatically excludes all branches in the phylogeny that are unrelated to the evolution of the convergent trait (dotted branches in **Fig. 1A**). This means that the model learning is directly focused on the molecular evolutionary changes between trait-positive and trait-negative species (solid blue and red branches, respectively). If there are multiple species in some trait-positive and trait-negative clades, different combinations of training sets may be used to build separate genetic models followed by model averaging (see *Material and Methods*). ESL-PSC analysis produces a list of proteins included in the genetic model, the estimated relative importance of each locus, and an equation to predict the presence/absence of the trait in a species based on its genetic sequences. Species not used for training for a given model can be utilized for testing the model.

## Genetic Models for Convergent Acquisition of C4 Photosynthesis

We applied ESL-PSC to build genetic models of photosynthesis evolution using a 64-species alignment of 67 chloroplast proteins^22^ (see *Material and Methods*). Many of these grass species have convergently evolved the C4 photosynthetic pathway for carbon concentration^24,25^, while others have retained the ancestral C3 photosynthetic pathway. Previous studies of the genetic basis of C4 evolution have found convergent amino acid substitutions in Ribulose-1,5-bisphosphate carboxylase-oxygenase (RuBisCo) to be strongly associated with C4 evolution, but Casola and Li^22^ have recently suggested the involvement of other chloroplast proteins as well. However, the extent to which chloroplast proteins other than RuBisCo represent a predictable and common evolutionary basis of C4 evolution remains uncertain.

There are six clades in the molecular phylogeny that contain sibling species of both C4 and C3 phenotypes (**Fig. 2**), which yielded six pairs of species satisfying the PSC design. Each pair contained a species with C4 photosynthesis and its most closely related species with C3 photosynthesis. Because some clades contain multiple candidate trait-positive (C4) and trait-negative (C3) species, we selected the species with the least missing data in the sequence alignment in our first analysis (solid lines in **Fig. 2**). The lengths of individual protein sequence alignments varied from 30 to 1,528 amino acids, with a total of 16,362 positions in 67 chloroplast proteins^22^.

**Figure 2.**
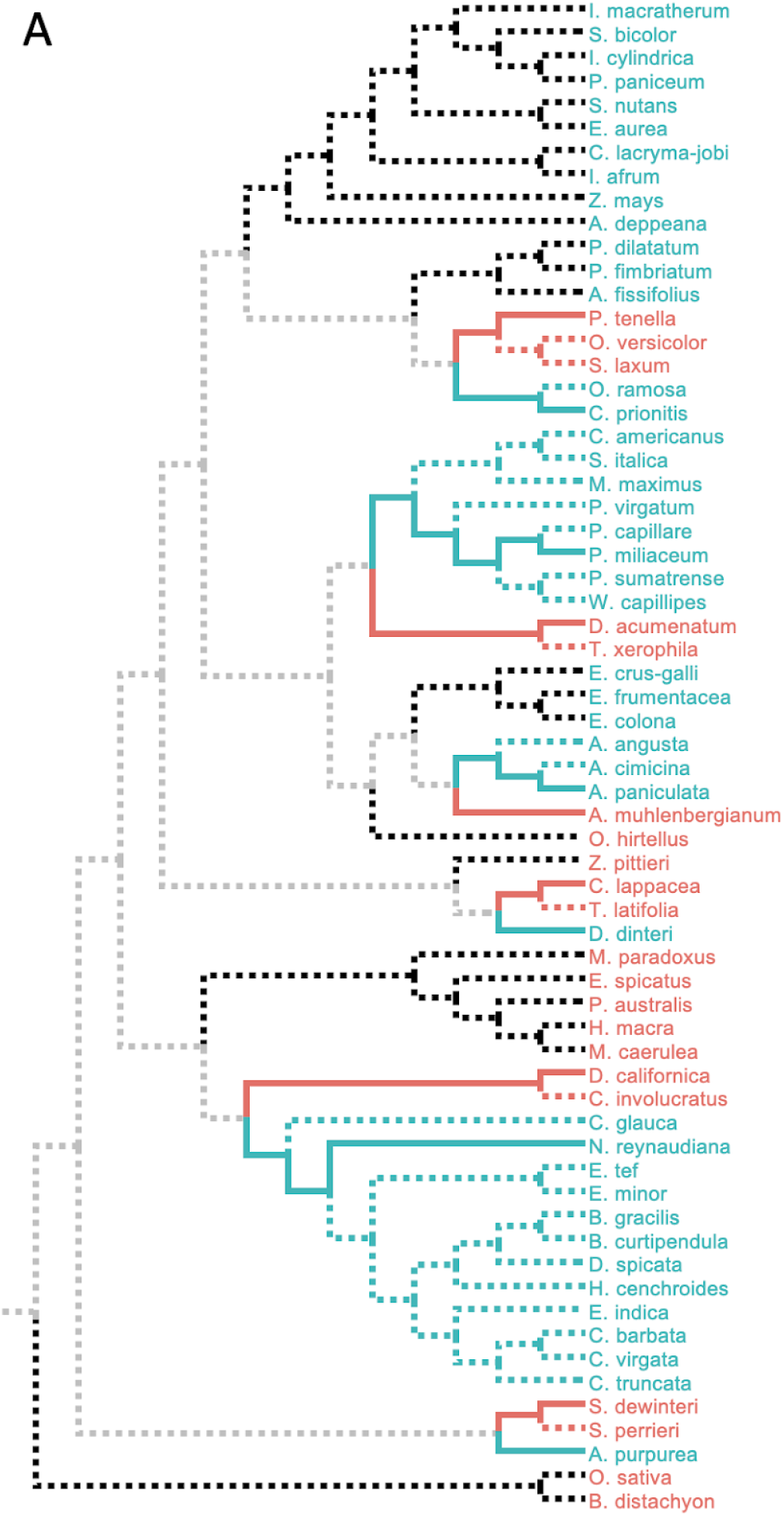
ESL-PSC modeling of convergent acquisition of C4 photosynthesis. **A. Experimental design.** An evolutionary tree of 64 grass species based on the phylogeny in Casola and Li ^22^. From the 64 available species, 6 pairs of trait-positive (C4) and trait-negative (C3) species were chosen according to the PSC approach. Where multiple species met the topological requirements for a contrast pair, we selected the two species that were closest in the evolutionary distance and that had the fewest gaps in the alignments. Selected species are shown as solid line branches, and all other branches are depicted as dashed lines. Solid lines begin at the internal node that represents the common ancestor of each pair, and the black (C4) and red (C3) branches represent the unshared ancestry of each selected species. Thus substitutions on these branches can be included in ESL-PSC modeling. Blue (C4) and red (C3) dashed lines represent alternative sibling species of the selected species. Black dashed branches represent clades that are evolutionarily independent of the contrast pairs. These include both C4 and C3 species. Gray branches represent the evolutionary history that is shared equally by selected C4 and C3 species, which we expect to cancel out automatically in the modeling process.

In ESL-PSC analysis, sparsity penalties must be specified for the inclusion of sites and proteins in the genetic model built using LASSO. These penalties dictate the number of proteins and sites allowed in the genetic model^8^. We used a series of penalties and compared resulting genetic models by using a newly developed Model Fit Score (MFS), which is analogous to the Brier score in logistic regression (see Methods). The genetic model with the best MFS contained included RuBisCo, consistent with previous experimental and analytical knowledge^20,22,26,27^. This model correctly assigned all six C4 and six C3 species used to train the model and correctly predicted 97% of the other C4 species in this dataset (36 of 37) and 100% of C3 species (15 of 15) for a balanced accuracy of 98.5%. An ensemble of genetic models with similar MFS scores (**Fig. 3**) also performed equally well (**Fig. 4A**).

**Figure 3.**
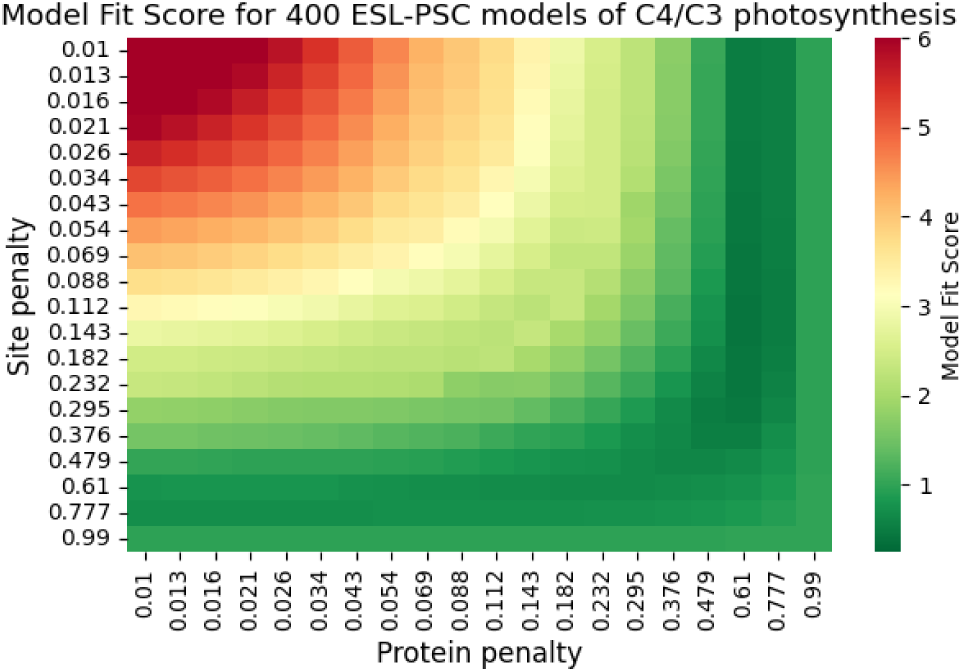
Heat map of Model Fit Scores. 20 values for each inclusion penalty (site and protein) were sampled from a logspace ranging from 1-99% of the maximum non-trivial penalty. A higher MFS suggests a higher risk of overfitting. Models with the best (lowest) 5% of MFS are included in predictive ensembles (Fig. 4, 5).

**Figure 4.**
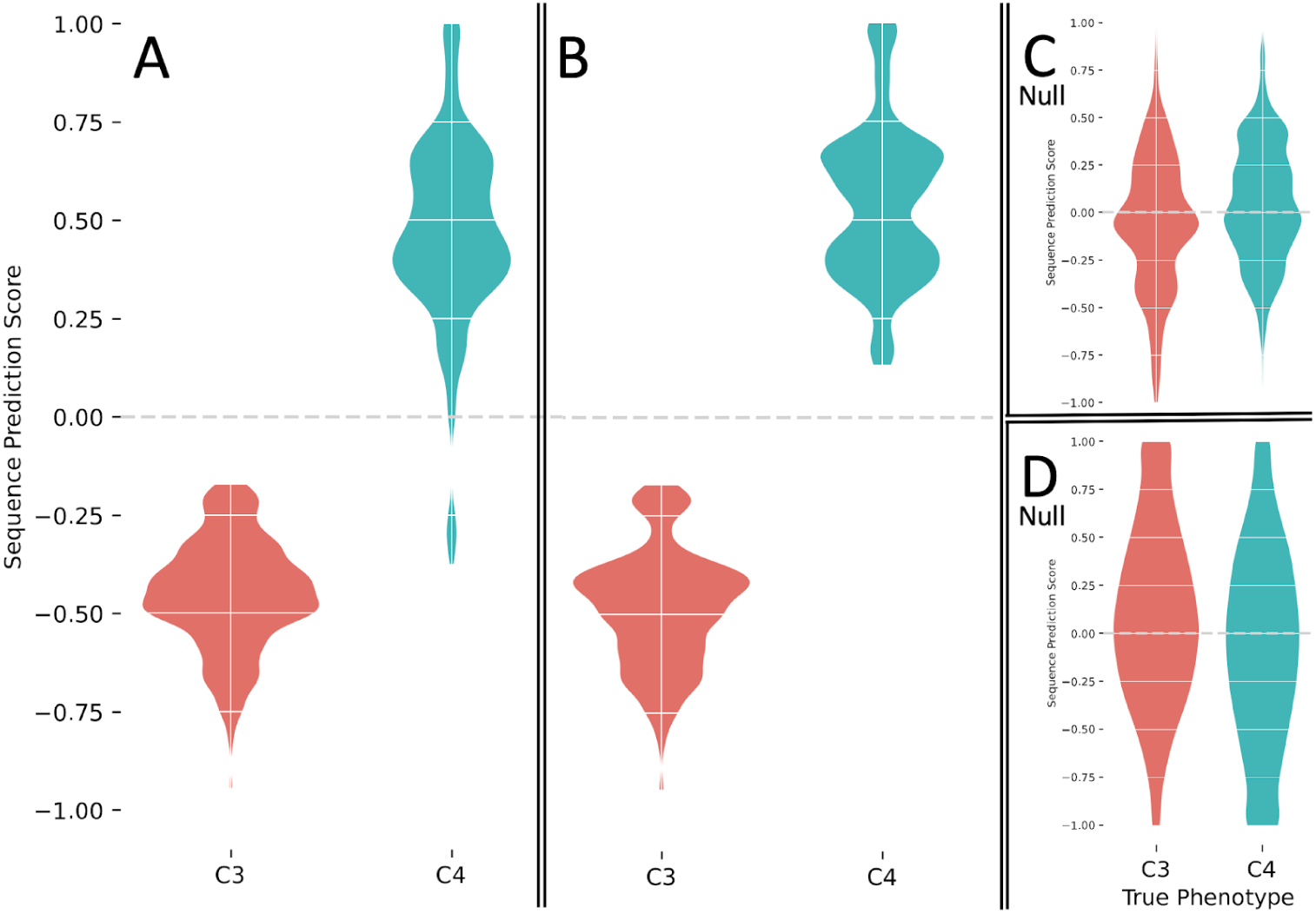
Predictive ability of ESL-PSC genetic models of C4/C3 photosynthesis. **A-D** Sequence prediction scores (SPS) from model ensembles are shown for known C4 (blue) and C3 (red) species in kernel density estimation plots. Negative SPS indicates a prediction of the C3 phenotype (trait-negative), and positive SPS indicates a prediction of the C4 phenotype. Predictions shown are for all species (A), species in clades independent of the clades contributing species for model training (B). Response-flipped null ESL-PSC models of C4/C3 photosynthesis (C). Null models were constructed by flipping the phenotype response values of 3 out of 6 of the input contrast pairs. This was done for all 10 distinct combinations of 3 out of 6 contrast pairs, and all model predictions were aggregated. SPSs from the best 5% of models by MFS are included. Pair-randomized null ESL-PSC models of C4/C3 photosynthesis (D). Null models were constructed by randomly flipping or not flipping the residues between each species contrast pair at every variable residue in the MSA. For each of the 25 alternative PSC input species combinations, randomized pair-flipped alignments were generated, and model ensembles were produced for each. Aggregated predictions are shown.

The best MFS model was found to be equally accurate in predicting C4 species that are siblings of those used in the training set, which suggests that multiple C4 species within a clade inherited the trait from a common ancestor. This is consistent with the parsimonious reconstruction of independent C4 trait evolution^28^. For this reason, genetic models built using different species combinations were also highly accurate (96%, **Fig. 5B**). The best MSF models were also highly predictive of the C4/C3 status of species from independent clades (black dotted branches in **Fig. 2**) that did not contribute any species for training the model (100% accuracy; **Fig. 4B**). This result suggests that many of the same substitutions contributed to C4 evolution independently.

**Figure 5.**
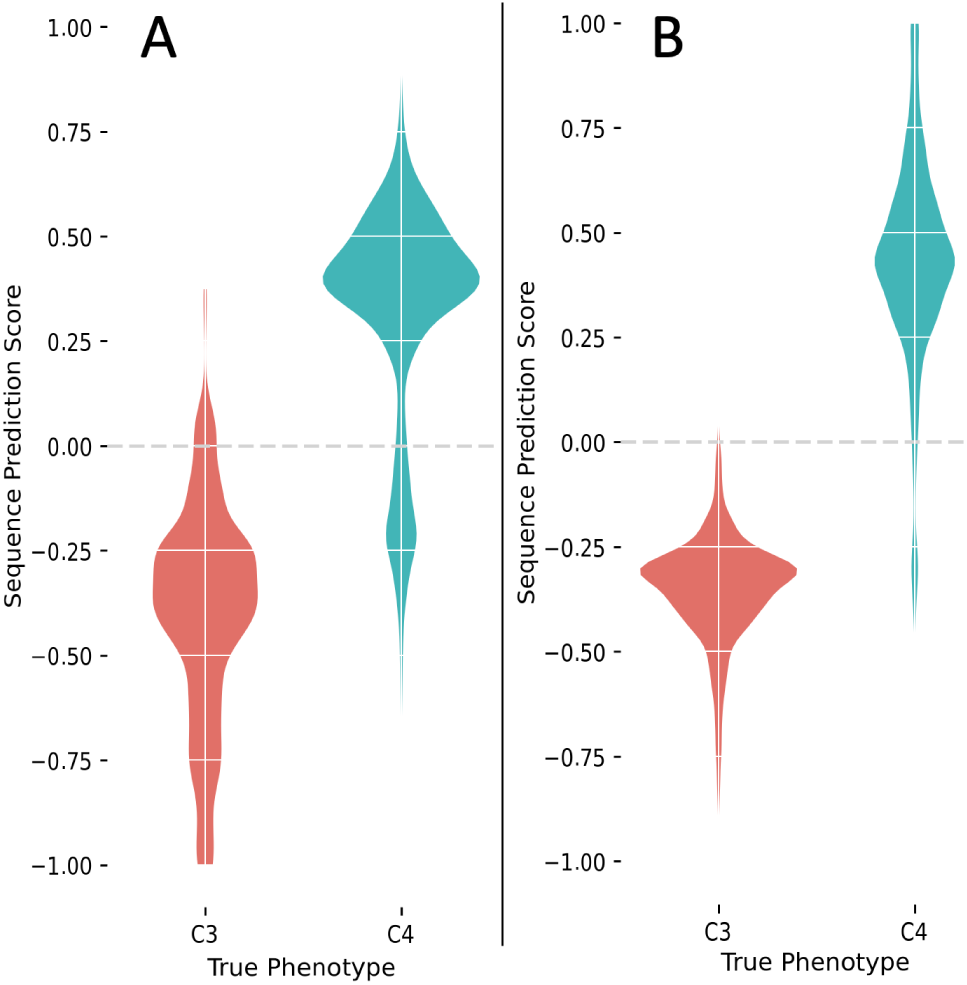
Alternative models. **A:** Predictions from models developed without the inclusion of RuBisCo are shown for independent species. **B:** Alternative PSC combinations. 100 alternative species combinations of PSC pairs were generated, and ensemble models were constructed as above. Predictions were aggregated for only the independent clades (black branches in Fig. 2). SPSs from the best 5% of models by MFS are shown from the aggregate of all ensemble models.

In addition, we found that evolutionarily-naive machine learning, which did not use the PSC design, could only achieve 64% accuracy in correctly identifying C4 species in the independent clades (black branches in **Fig. 2**). In this experiment, we conducted a direct comparison by selecting 100 input sets of six C4 and six C3 species from among the siblings of the PSC species, but without respecting the PSC design. For these “naive” models, the prediction accuracy fell considerably. In particular, the average true positive rate (TPR), a measure of the ability of the model to recognize C4 species on the basis of information in convergent sites, was only 64% over all of these ensembles compared with 94% for the ensembles built using the PSC approach (**Fig. 4B**). This reduction in accuracy reflects the fact that non-PSC models may incorporate not only sites whose residues are correlated with the phenotype due to convergent evolution but also sites correlated with the phenotype purely due to shared ancestry within the inputs. The latter type of sites carries no information relevant to the prediction of phenotype in clades whose trait-positive species have acquired the trait independently. This result establishes that our PSC design can produce much better genotype-phenotype models than naive machine learning.

Studies of convergence in C4 have focused heavily on RuBisCo, the most abundant enzyme, which has multiple sites of convergent amino acid substitutions in multiple different lineages of plants^20–22,26,29^. However, we tested the hypothesis that other chloroplast proteins also contributed to C4 evolution by building ESL-PSC models excluding RuBisCo and testing model accuracy in predicting the presence of C4. The RuBisCo-free models had 89% accuracy, suggesting that the convergent basis of the C4 trait extends to other chloroplast genes (**Fig. 5a**). Interestingly, these models correctly predicted C4 photosynthesis in *Alloteropsis angusta*, which was the only false negative for the model containing RuBisCo. *A. angusta* is known to have undergone a C3 to C4 transition independently from the other members of its own genus, including *A. paniculata*^30^. We found *A. angusta* to be lacking key amino acid substitutions in RuBisCo that are highly diagnostic of other C4 species. Therefore, chloroplast proteins other than RuBisCo have likely contributed significantly to C4 evolution in this case, and more generally. While Casola and Li^22^ hinted at such a possibility, their statistical analyses using a convergence counting approach did not find a significant excess of convergent substitutions in C4 species as compared to the background C3 species. Therefore, the ESL-PSC framework provided a powerful new way to investigate the genetics of convergent traits and test hypotheses that have not been possible until now.

### Convergent Evolution of Echolocation

The independent acquisition of echolocation in bats and whales is among the most well-studied cases of convergent molecular and trait evolution. We selected the microbat *Myotis lucifugus* and the bottlenose dolphin *Tursiops truncatus* as trait-positive species (echolocators) because previous studies involving exome-scale searches for convergence in echolocating mammals have often focused on the comparison of microbats and toothed whales^2,3,9,31^. In the PSC design, we selected a non-echolocating sister species *Pteropus vampyrus* (megabat) for echolocating *Myotis lucifugus* and non-echolocating *Ovis aries* (sheep) for echolocating bottlenose dolphin *Tursiops truncatus* (**Fig. 6**; see *Methods*). We retrieved 14,509 protein alignments from the OrthoMaM database of orthologous protein-coding sequences for mammalian genomes^32^.

**Figure 6.**
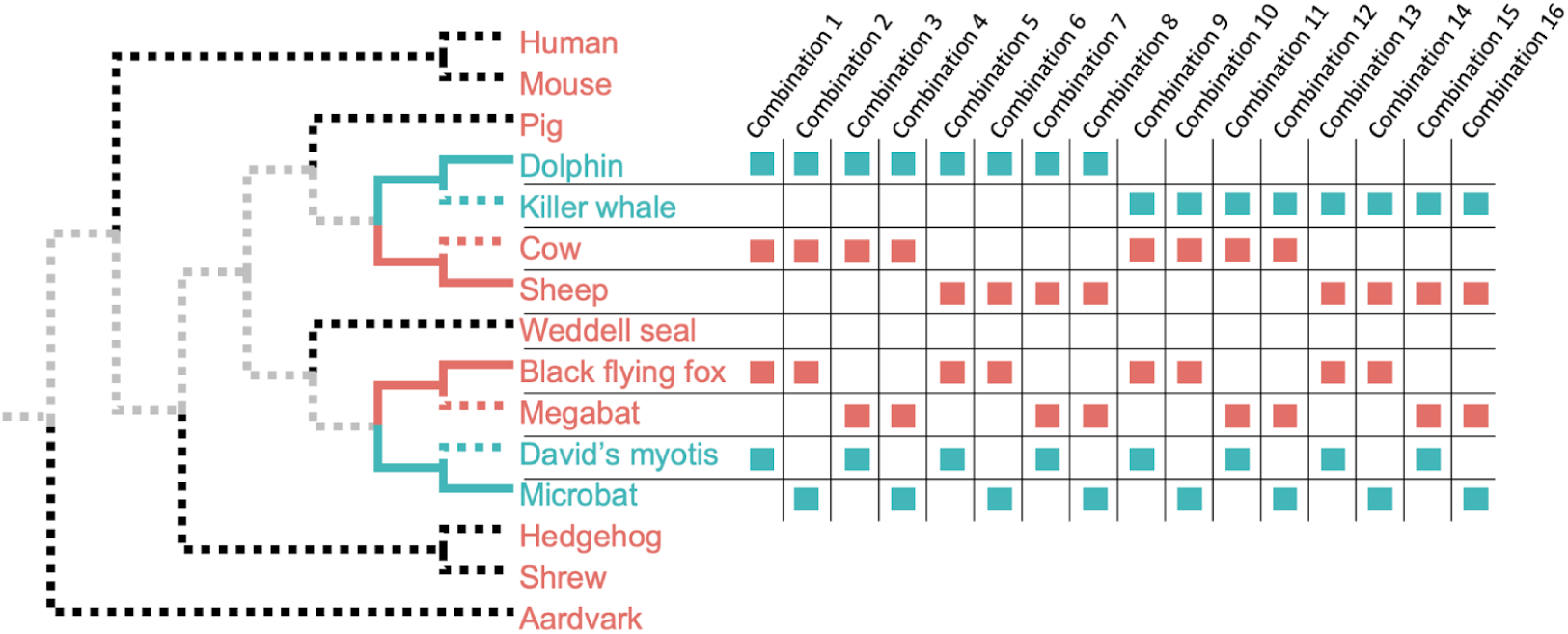
Echolocation analysis. Echolocation evolved twice in mammals in our dataset. Therefore two contrast pairs can be constructed (solid blue branches, echolocating; solid red branches, non-echolocating). A series of 15 comparable sets of input pairs can be constructed by using alternative species (dashed blue and red sibling species) in all possible combinations. Species not included in the contrast pairs do not affect the analysis (black dashed branches). Shared ancestry is canceled out (gray branches).

Because there were only two clades, and thus only two species pairs, we made inferences from a collection of ESL models obtained using a range of sparsity penalties (see *Methods*) and species combinations (**Fig. 6**). The collection of genetic models was then used to generate a ranked list of candidate proteins associated with convergent evolution (**Table S1**). Among the highest-ranked proteins, many were those previously characterized to have signatures of molecular convergence in echolocators, including, Prestin (SLC26a5), TMC1, PJVK (DFNB59), CDH23, CASQ1, and CABP2^3,17,18,33–35^. In some cases, specific amino acid sites within these proteins have been implicated in conferring the functional changes necessary for the echolocation phenotype, revealed by laboratory assays where mutations to residues found in echolocating species were observed to alter protein function in a manner consistent with echolocation^3,19^.

We generated multiple-tests adjusted *P*-values to gauge the functional enrichment in the top-ranking proteins included in the genetic models. We tested for ∼20,000 biological processes and phenotypes (see *Methods*) and found the top 100 proteins to be highly enriched for the “sensory perception of sound” genes (GO:0007605) with an adjusted *P*-value < 10^−4^ (**Table 1**). This is an improvement in the statistical significance of more than two orders of magnitude compared to the best previous findings of this term (adjusted *P* = 0.049) in FDR-corrected analyses^4,31^. Our enrichment *P*-value was highly significant even for 50, 150, and 200 top proteins in the genetic models (*P* < 10^−3^), suggesting that our results are robust to the size of the gene list analyzed.

**Table 1.** Ontology term enrichments. Enrichment tests were performed for Gene, Phenotype, and Disease ontology terms for the top 100 highest-ranking trait proteins in our echolocation multiple species combination ensemble model integration analysis. In each figure, the 10 ontology terms with the lowest p-values are shown from each enrichment analysis.

| Term | P-value | Adjusted P-value |
| --- | --- | --- |
| **Go Biological Process** |  |  |
| sensory perception of sound (GO:0007605) | $< 1 \times 10^{-7}$ | $< 1 \times 10^{-4}$ |
| sensory perception of mechanical stimulus (GO:0050954) | $< 1 \times 10^{-6}$ | $< 1 \times 10^{-3}$ |
| **MGI Mammalian Phenotype Level 4** |  |  |
| cochlear inner hair cell degeneration MP:0004398 | $< 1 \times 10^{-6}$ | $< 1 \times 10^{-3}$ |
| cochlear ganglion degeneration MP:0002857 | $< 1 \times 10^{-6}$ | $< 1 \times 10^{-3}$ |
| head tossing MP:0005307 | $< 1 \times 10^{-6}$ | $< 1 \times 10^{-3}$ |
| increased or absent threshold for auditory brainstem response MP:0011967 | $< 1 \times 10^{-6}$ | $< 1 \times 10^{-3}$ |
| organ of Corti degeneration MP:0000043 | $< 1 \times 10^{-5}$ | $< 1 \times 10^{-3}$ |
| deafness MP:0001967 | $< 1 \times 10^{-4}$ | $< 1 \times 10^{-2}$ |
| **DisGeNet** |  |  |
| Sensorineural hearing loss, bilateral | $< 1 \times 10^{-8}$ | $< 1 \times 10^{-5}$ |
| Nonsyndromic Deafness | $< 1 \times 10^{-4}$ | $< 1 \times 10^{-1}$ |

This top-100 gene list was also significantly enriched (adjusted *P* < 1×10^−6^) for many Phenotype Ontology (PO) terms directly related to hearing and sound perception such as “cochlear inner hair cell degeneration” (MP:0004398), “increased or absent threshold for auditory brainstem response” (MP:0011967), “cochlear ganglion degeneration” (MP:0002857), and “increased or absent threshold for auditory brainstem response” (MP:0011967) (**Table 1**). We also found a highly significant enrichment (adjusted *P* < 4.5×10^−3^) for the top-level mammalian PO term “hearing/vestibular/ear phenotype” (MP:0005377).

As a control, we built null genetic models in which one of the two contrast pairs had its trait status reversed, such that the echolocating dolphin and non-echolocating large flying fox were treated as sharing a convergent trait, while the other two species were treated as paired contrast partners. This configuration has the property that both the shared phylogenetic signal and any shared convergent trait signal from the genuine trait of echolocation are canceled out. Then, we applied GO and PO enrichment to the top 100 genes in the ESL-PSC models as above. None of the terms in **Table 1** received significant enrichment (adjusted *P* < 0.05), as expected of the null model. A recent study found that the analysis of synonymous variation can help detect data contamination and other types of error ^36^, so we developed another null test of ESL-PSC by analyzing only fourfold degenerate sites expected to evolve largely neutrally in mammals. No significant enrichment was found for any of the relevant ontology categories.

Overall, highly significant probabilities for the enrichment of hearing-related ontology terms suggest that machine learning detects a strong signal of convergence in hearing-related proteins in echolocators. This is the first demonstration of a multiple test-adjusted highly significant signal for sound perception in a genome-wide comparative analysis of echolocation.

## DISCUSSION

Discovery of genotype-phenotype relationships is of central importance in functional and evolutionary genomics. Repeated evolution of the same trait in species of independent clades offers an opportunity to reveal the genetic architecture shared by these independent trait evolutions. We have presented a novel comparative genomics approach using machine learning (ESL-PSC), informed by molecular phylogenies, to infer quantitative genetic models of trait convergences. The application of ESL-PSC to two distinct, previously well-investigated examples establishes that there is a significant commonality in the genetic basis of trait evolution among species in independent lineages.

A high predictive ability of ESL-PSC was found for correctly classifying species with and without C4 photosynthesis in grass clades not involved in training the model (**Fig. 4A, B**). Classical molecular evolutionary methods do not commonly afford this type of quantitative prediction. The high accuracy of genetic models of C4 trait evolution in which the well-studied convergent protein RuBisCo was excluded is suggestive of the potential role of additional chloroplast proteins in the convergent gain of C4 photosynthesis. These analyses also showed that not all species with convergent traits harbor the same substitutions in the sites included in genetic modes. In fact, no more than four out of six C4 species shared the same amino acid residue in the sites selected during ESL model building. Therefore, ESL model building can automatically extract relevant information from incomplete molecular convergence correlated with the trait convergence, obviating the need to use *ad hoc* cut-offs and subsetting data by evolutionary conservation^2,3,5,7,37^. This makes ESL-PSC convergent evolution analyses less subjective and more reproducible than other approaches.

ESL-PSC also identified genes involved in the convergent acquisition of echolocation in mammals. The list of top genes in ESL models was found to be highly enriched for GO and PO categories involved in auditory processes at FDR-corrected *P*-values that were more significant than previously reported, implying that the machine learning approach to building genetic models can be significantly more powerful than previous approaches. While validation of ESL-PSC derived from the enrichment of functional categories is arguably circumstantial, direct experimental approaches are beyond the scope of this investigation. Further support may be found by assessing the potential functional relevance of the selected genes to determine whether mutations in them cause diseases due to relevant functional disruptions. In the analysis of Disease Ontology categories, we found a hearing-related “Sensorineural hearing loss, bilateral” term to be highly enriched in the top genes (adjusted *P* < 10^−5^). Many other terms related to deafness contained a significantly greater than expected number of genes (**Table 1**). No previous study has reported such an enrichment.

ESL-PSC appears to extract commonalities of the genetic basis of trait convergences more effectively than other approaches. However, we note that species-specific evolutionary substitutions may also be involved in the evolution of convergent traits. These are not the target of the ESL approach and will not be included in the genetic model. Also, molecular convergences in the non-coding sequences as well as regulatory innovations may be involved in the evolution of convergent traits some of which may be analyzed by their simultaneous analysis in the ESL-PSC framework. We plan to pursue them in the future.

We expect ESL-PSC to be useful as a comparative genomics tool for uncovering common genetic elements involved in the evolution of traits shared between species. We envision that ESL-PSC will be applied to first generate a candidate gene and site list, which can be followed by a series of hypothesis tests regarding the commonality of the genetic basis of trait convergences. These analyses will be extremely fast, as ESL-PSC took only minutes in most of our data analyses. These results can then be followed up by conducting traditional molecular evolutionary analyses and functional genomic experiments to identify selective processes at play.

## Acknowledgments

The authors would like to thank Drs. Alessandra Lamarca, Jack Craig, and Sayaka Miura for reading the manuscript and providing many helpful suggestions. This work was supported by research grants from the National Institutes of Health to SK (R35GM139540-03) and a fellowship to JA from Temple University.

## Author information

### Contributions

S.K. conceived the idea and developed the initial method; J.A. and S.S. refined and extended the method; M.S., S.S, J.A., and R.P. implemented the method; J.A. and R.P. conducted the data analyses; J.A., S.K., and G.G. wrote the manuscript; all authors contributed to intellectual discussions about the method and results and co-wrote the manuscript.

## Methods

### Genomic alignment data retrieval and processing

Alignments of chloroplast genes were retrieved from the supplemental data in ref.^22^. We generated translated amino acid sequences from the provided nucleic acid alignments for ESL-PSC analyses. The OrthoMaM data set^32^ of mammalian one-to-one orthologous protein sequence alignments was downloaded from https://orthomam.mbb.cnrs.fr/. Following previous studies in which exome-scale scans for convergence in echolocating mammals were performed, we analyzed echolocation in microbats and toothed whales^2–4,9,31^ and used megabats and artiodactyls as non-echolocating sister taxa^2,5,6,9^. In their ESL-PSC analysis, we excluded sites containing missing data or alignment gaps in individual training sets. All multiple sequence alignments (MSAs) were one-hot encoded^8^, which transforms it into a numerical format that is required by the model-building algorithm. The presence of the convergent trait was represented numerically by +1 and its absence by −1.

### Building Genetic Models

ESL-PSC uses the Least Absolute Shrinkage and Selection Operator (LASSO)^23^ logistic regression, in which coefficients are chosen to minimize a combination of the difference between observed and predicted response values of the input species (the logistic loss). It uses an inclusion penalty term that scales with the sum of the absolute values of the model coefficients and, therefore, induces sparsity^8^. We use bilevel sparsity in which separate penalties are applied for the inclusion of sites and groups of sites (e.g., proteins). The loss function is minimized by gradient descent ^38^, which is re-implemented in the myESL software package used for ESL-PSC implementation (https://github.com/kumarlabgit/ESL-PSC). We estimate a new Model Fit Score (MFS) for a given genetic model, which is the root mean squared difference between the input trait value (+1 and −1) and predicted trait values for all species used for training the model. The best-fit genetic models have the lowest MFS value, i.e., the input and output of the genetic model are the most concordant. This is needed because optimal inclusion penalties are not known beforehand in LASSO. So, the genetic model with the best MFS is chosen.

In our analysis, the size of the penalty for each protein (group of sites) was globally controlled by the inclusion penalties, but can also vary for each individual group depending on its composition. Group penalties in applications of the LASSO method are typically based on the square root of the number of columns belonging to the group in dataset^39^. Applying this system produced models in which proteins with fewer variable sites and lower total entropy were penalized more than those with many variable sites, in the exome-wide analysis. However, highly conserved proteins containing even a few variable sites can be important. Therefore, we devised a penalty function for each protein in which the group penalty scales linearly with the number of variable sites plus a constant equal to the median number of variable sites across the proteins in the dataset (excluding fully invariant proteins). This function was effective for both small-scale (chloroplast exome) and large-scale (mammalian proteome) analyses.

### Predictive Model Ensembles

Models with similar MFS scores were combined to form ensembles of models for predictions. For all model ensembles, we used a range of group and site inclusion penalty values from 1%-99% of the maximum penalty that can be applied before a trivial solution in which all model feature weights are set to 0 is obtained. The inclusion penalty values were taken from a logspace over this range. Unless specified, we selected genetic models with the best MFS or those with the top-5% MFS values.

### Building the Candidate protein list

We estimate the Group Sparsity Score (GSS) for every selected protein in every model over all inclusion penalty combinations. GSS is the sum of absolute values of regression coefficients for all the selected positions in the given protein^8^. The higher the GSS, the greater their importance. Proteins not included in the genetic model receive GSS = 0. For every candidate gene, their overall rank is the best rank (according to their GSS) they receive in any of the genetic models, with equally ranked proteins being further ordered according to the maximum GSS they attained in any model. This yields an ordered list of proteins whose convergent sites stand out compared with the rest of the proteome in number, proportion, and strength of the concordance of their convergent site patterns with the species phenotypes, without privileging any one of those considerations.

When each of the input species has at least one sibling species that share its phenotype for the trait being studied, then different combinations of these allowable input species can be used interchangeably, and models over all inclusion penalty combinations can be built for each of the species combinations. The output candidate convergent proteins are then ranked by the number of species combinations for which they received non-zero GSS scores in at least one model, with ties being resolved by the number of species combinations in which the proteins were ranked in the top 1%, followed by the highest ever rank and highest ever GSS obtained.

### Ontology analysis

Ontology enrichment testing was performed using Enrichr^40^, and *P*-values were adjusted for multiple testing. Gene ontologies were obtained from GO^41^. We tested for the biological process GO ontologies using the GO_Biological_Process_2021 set in Enrichr (6,036 terms). Phenotype ontologies were derived from MGI^42^. Enrichr provides PO testing using a trimmed version of the MGI phenotype vocabulary. Which excludes the top three levels of PO terms (4,601 terms). Disease ontologies were derived from DisGeNet (9,828 terms)^43^. To determine enrichment and overlapping genes for the top-level PO term “hearing/ vestibular/ ear phenotype” (MP:0005377), we used the MouseMine ^44^ ontology testing tool and the Benjamini-Hochberg adjustment to obtain a multiple testing adjusted P-value. By common convention, enrichments were only considered valid if accounted for by an overlap of at least 5 genes. Phenotype ontology terms were retrieved from the Mouse Genome Informatics mammalian phenotype vocabulary, and gene lists associated with phenotype ontology terms were generated from the Mouse/Human Orthology with Phenotype Annotations (downloaded from http://www.informatics.jax.org/downloads/reports/index.html#pheno). For gene enrichment analyses, we found that it was unnecessary to use ensembles of 400 models (20 values for each inclusion penalty) because the gene ranks are based on the maximum model weights which do not change significantly when using a denser grid search over the space of inclusion penalty. Results shown here were based on ensembles using 4 values of each inclusion penalty (16 models) in each ensemble for each species combination.

### Null Genetic Model Ensembles

There are a number of different ways to test the genetic models produced by machine learning. We built null genetic models by reversing trait designations of a subset of training data such that both the shared evolutionary history and shared basis of the convergent trait between trait-positive species were canceled out (**Fig. 3C**). For an even number 2*n* of input species contrast pairs, the largest scrambling of the input phenotype designations is achieved by flipping n pairs. There are ½^2*n*^C*_n_* possible distinct null configurations. For a small *n*, it is possible to generate and combine all null predictions, but a random subset of possible null configurations can be sampled when *n* is large. Another type of null model can be constructed by randomly flipping (or not flipping) the residues between the two members of each contrast pair at each site (**Fig. 3D**). This preserves any phylogenetic relationships present in the alignment but, when averaging over a large number of such pair-randomized alignments, destroys the correlations that are due to convergence. Both of these null model experiments are expected to produce models whose prediction accuracy on test species not used in model building is comparable to random chance. Protein lists developed by using null genetic models are not expected to be enriched in any functional ontology terms beyond that expected by random chance alone.

## Data availability

Grass and mammalian protein sequence alignment data required to reproduce the analyses in this article can be found at: https://github.com/kumarlabgit/ESL-PSC.

## Code availability

A GitHub repository containing scripts and software used to perform the ESL-PSC analyses in this study is available at https://github.com/kumarlabgit/ESL-PSC.

## Extended data

**Supplementary Table 1:**
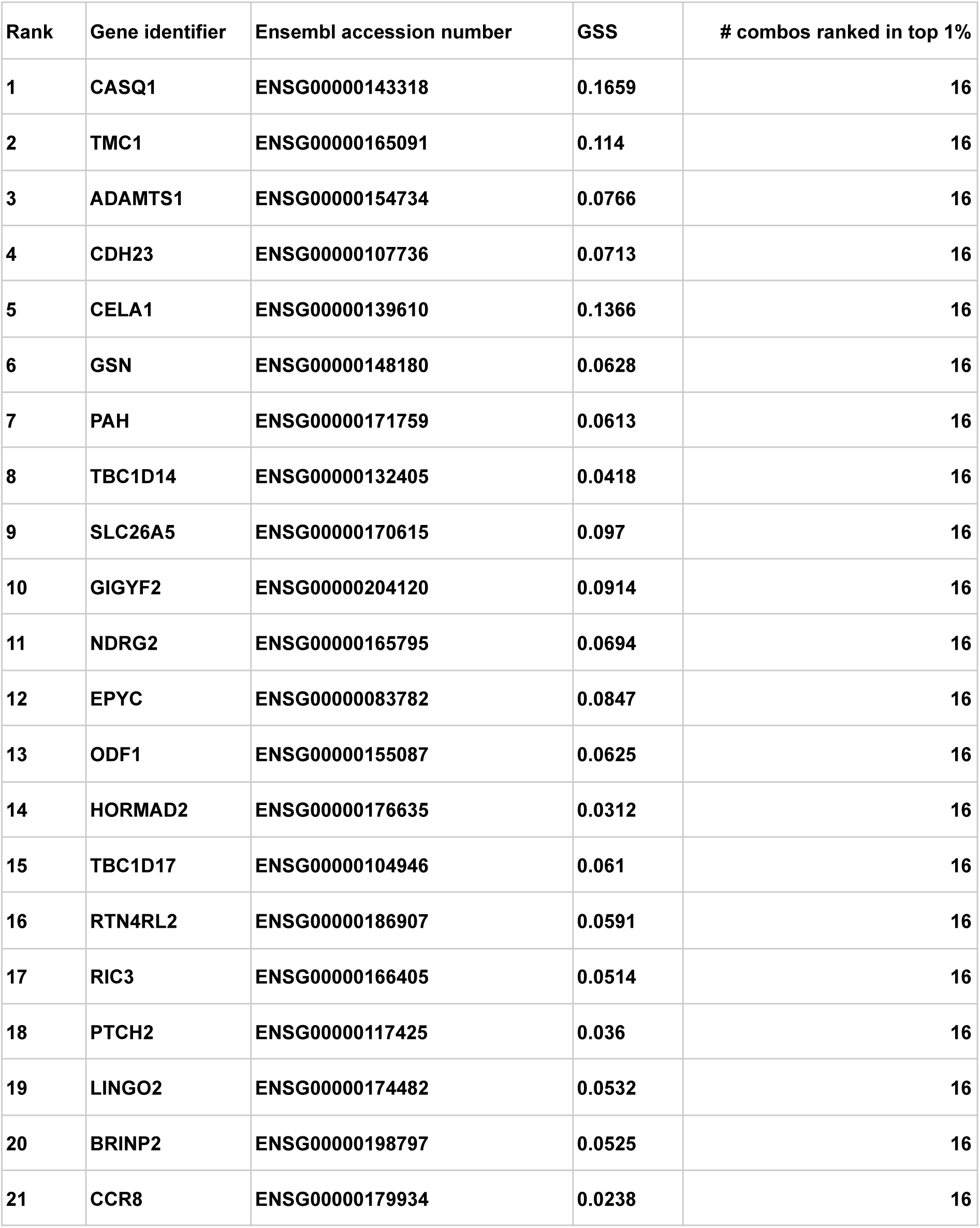
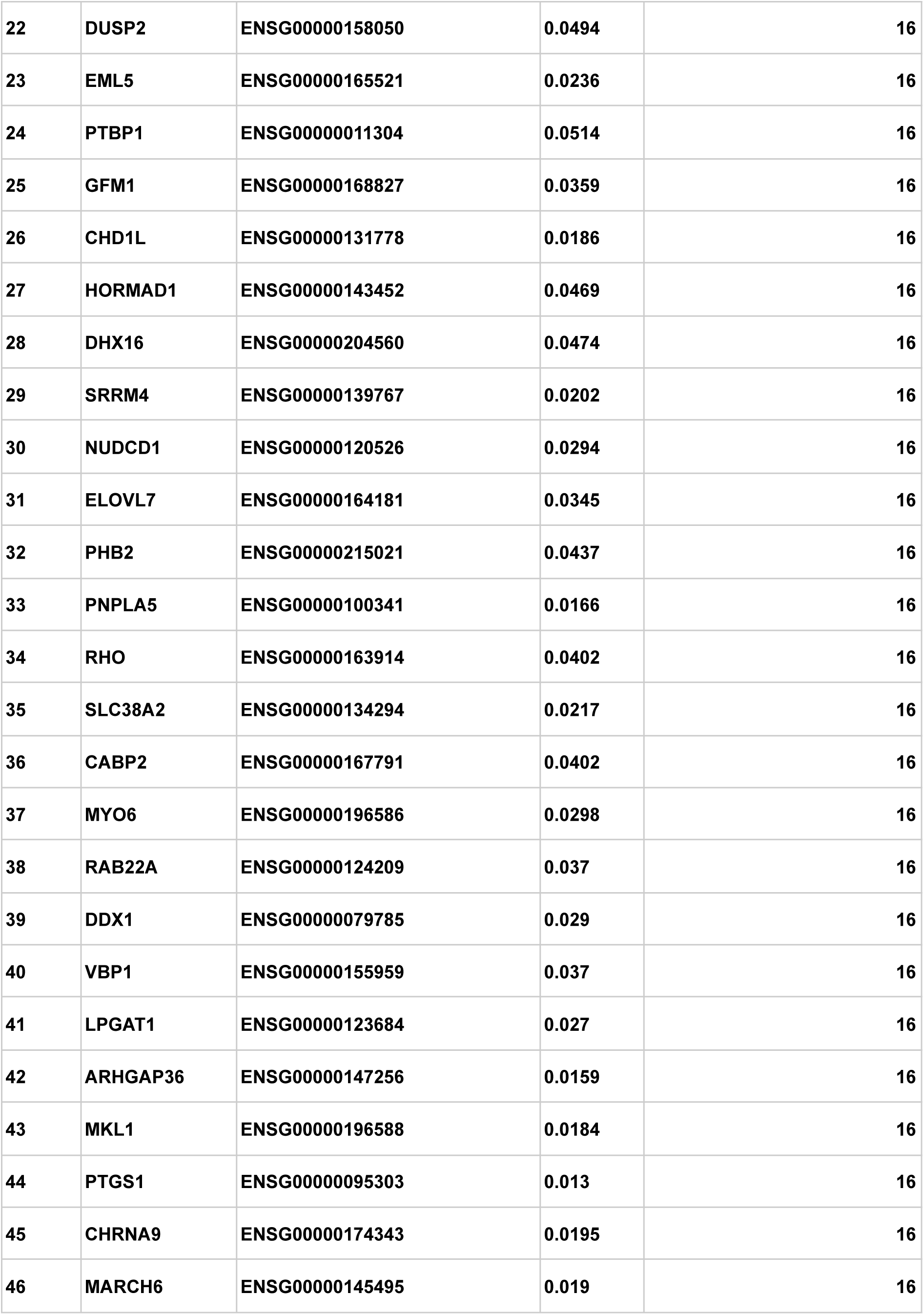
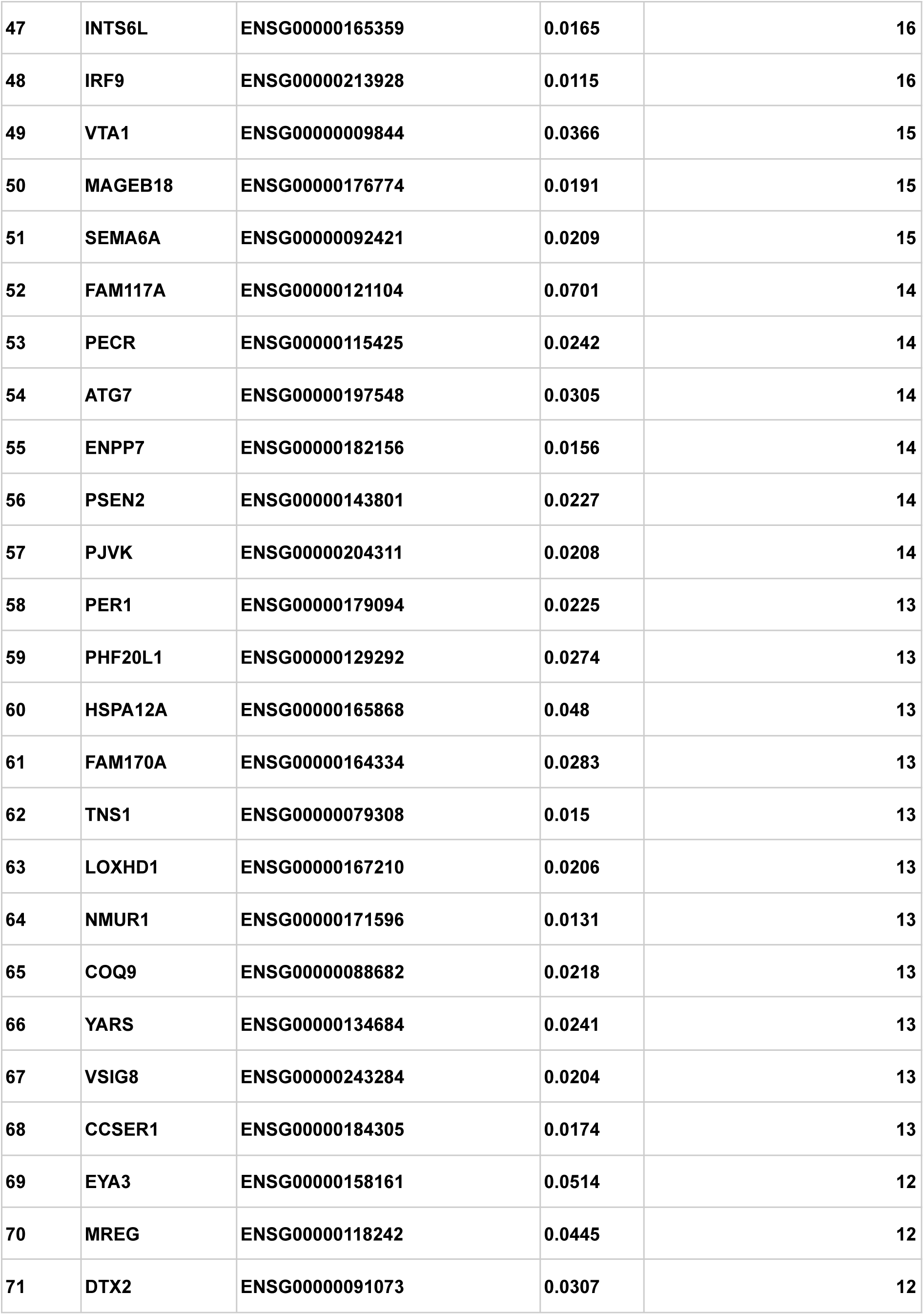
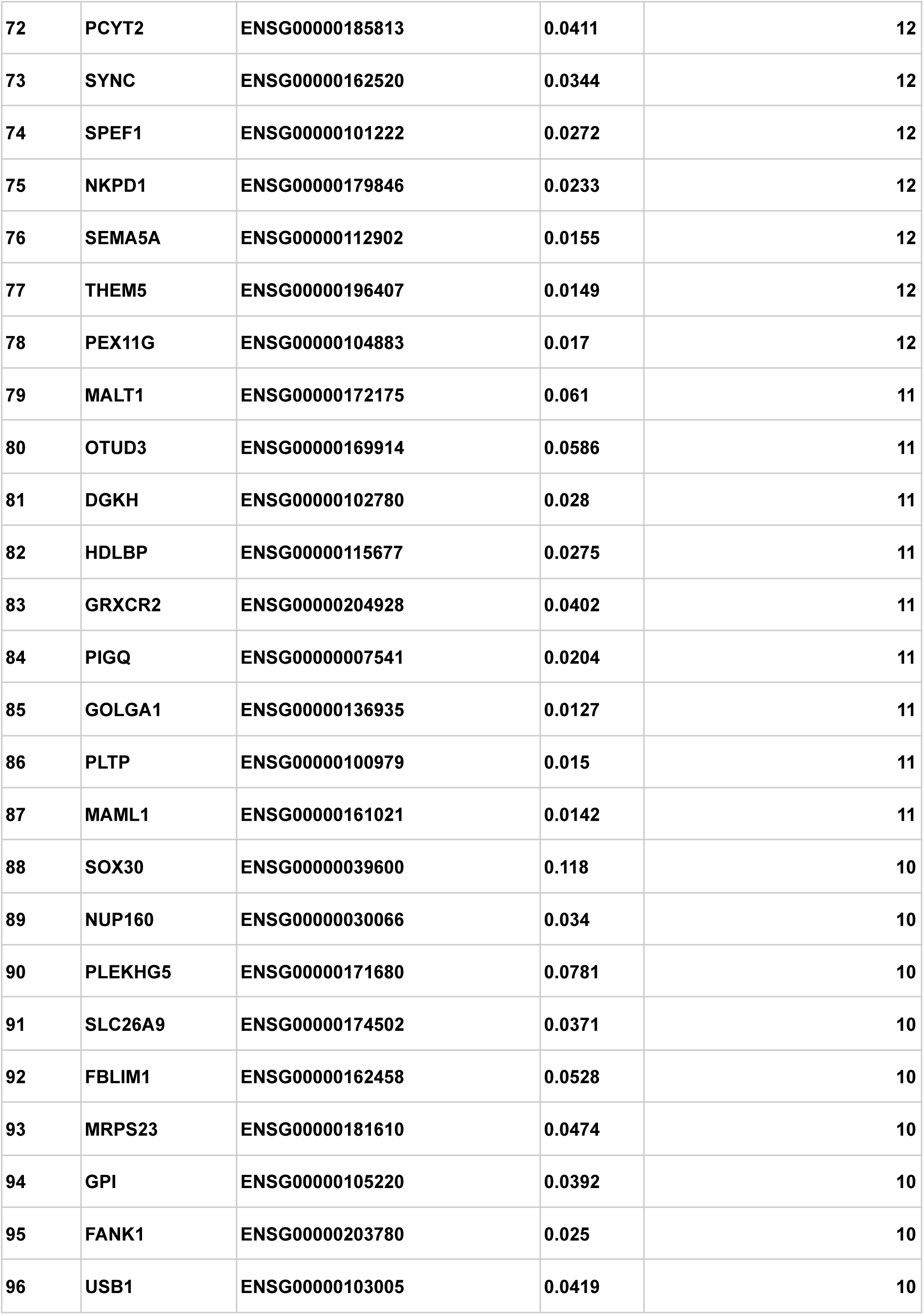
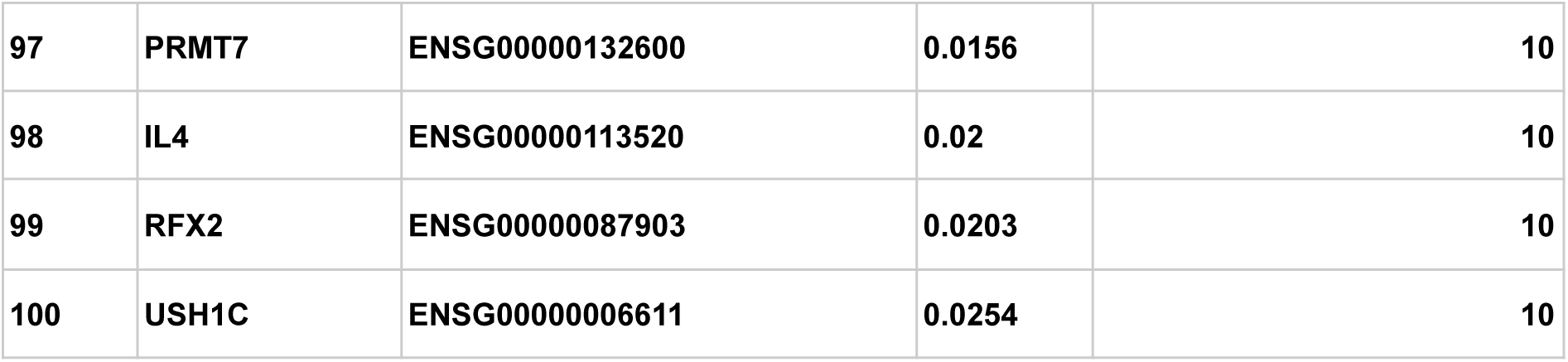
Echolocation ensemble model top genes.

